# Accurate Modeling of Brain Responses to Speech

**DOI:** 10.1101/509307

**Authors:** Daniel D.E. Wong, Giovanni M. Di Liberto, Alain de Cheveigné

## Abstract

Perceptual processes can be probed by fitting stimulus-response models that relate measured brain signals such as electroencephalography (EEG) to the stimuli that evoke them. These models have also found application for the control of devices such as hearing aids. The quality of the fit, as measured by correlation, classification, or information rate metrics, indicates the value of the model and the usefulness of the device. Models based on Canonical Correlation Analysis (CCA) achieve a quality of fit that surpasses that of commonly-used linear forward and backward models. Here, we show that their performance can be further improved using several techniques that capture the time-varying and context-dependent relationships within the data, including adaptive beamforming, CCA weight optimization, and recurrent neural networks that capture the time-varying and context-dependent relationships within the data. We demonstrate these results using a match-vs-mismatch classification paradigm, in which the classifier must decide which of two stimulus samples produced a given EEG response and which is a randomly chosen stimulus sample. This task captures the essential features of the more complex auditory attention decoding (AAD) task explored in many other studies.

## 1 Introduction

In experiments that record brain responses to stimulation, stimulus-response models are useful in providing insight into the components of the response. As these models can provide information about auditory attention, they have also been put forward for brain-computer interface (BCI) applications, such as the “cognitive” control of a hearing aid [Wronkiewicz et al., 2016]. Previous studies have used linear system identification techniques to either predict the response from the stimulus (forward model) or else infer the stimulus from the response (backward model) [Lalor and Foxe, 2010, Ding and Simon, 2012a,b, 2013, 2014]. In addition to these, a third form of model projects both stimulus and response into a common subspace via weight matrices obtained using Canonical Correlation Analysis (CCA) [Hotelling, 1936, Dmochowski et al., 2017, de Cheveigné et al., 2018]. As they are applicable to responses to arbitrary stimuli, they allow the research to move beyond the standard “evoked-response” paradigm that requires repeating the same short stimulus many times [Ross et al., 2010]. The quality of the model can be quantified by calculating the correlation coefficient between actual and predicted brain response (forward model), or between the actual and inferred stimulus (backward model), or between canonical correlate (CC) pairs (CCA). Higher correlation values indicate that the model better captures the relation between stimulus and response.

Alternatively, the quality of a model can be quantified on the basis of its performance in a classification task, in terms of discriminability (d-prime) or percent correct classification. This is particularly useful when developing a model for BCI applications where classification decisions are made based on short segments of data. In this paper, we use a simple “match-vs-mismatch” task based on the cortical response to a single speech stream [de Cheveigné et al., 2018], in which the classifier must decide whether a segment of EEG matches the segment of stimulus that evoked it, as opposed to some unrelated segment of the same stimulus. A good classification performance is taken to indicate that the model successfully captures the stimulus-response relationship.

Other studies have used the more complex Auditory Attention Decoding (AAD) task, in which a subject is presented with two concurrent stimulus streams (for example two voices speaking at the same time) and required to attend one stream or the other. The classifier attempts to identify which stream was the focus of the subject’s attention, given both stimulus streams and the EEG [Hillyard et al., 1973, Ding and Simon, 2012b, Mirkovic et al., 2015, 2016, O’Sullivan et al., 2015, Akram et al., 2016, O’Sullivan et al., 2017]. Our simpler task allows a more direct evaluation of the stimulus-response model that underlies both tasks.

A previous study from our group found that models based on CCA were superior to classic forward and backward models in terms of correlation, d-prime, and classification error rate [de Cheveigné et al., 2018]. Better performance was attributed to the ability of CCA to strip both stimulus and EEG of irrelevant dimensions, and to the fact that the multiple CCs allow multivariate classifiers to be deployed. In the aforementioned study, the various models were constrained to have the same number of free parameters so as to ensure a fair comparison between models. Here, we relax that constraint and introduce several new schemes to improve model quality. Arguably, models that give better performance more accurately capture the cortical representation of the stimulus, and good performance is also essential for applications. Each strategy is evaluated individually and in combination with others by comparison with a baseline (backward model or CCA).

Apart from the standard backward model, we test the following models and classification schemes (each coded by a letter): *CCA* (C), *maximizing component d-prime* (D), *adaptive beamforming* (B), *linear discriminant analysis* (L), *multilayer perceptron* (M), *simple recurrent layer* (S) and *gated recurrent unit* (G). Both *D* and *B* improve the computation of CCA components. *D* does this during training, and *B* does this during testing. *M, S* and *G* use a neural network architecture to improve the match-vs-mismatch classification over *L*.

## 2 Methods

### 2.1 Evaluation Dataset

The dataset used to evaluate canonical correlation analysis (CCA) performance was presented in [de Cheveigné et al., 2018] and published in [Di Liberto et al., 2015, Broderick et al., 2018a,b]. The speech stimulus was an audio book recording of the “Old Man and the Sea” recorded with a 44100 Hz sampling rate. The recording was divided into 32 segments lasting approximately 155s each. The stimulus was presented diotically over headphones to 8 subjects, while electroencephalography (EEG) data were recorded using a 128-channel Biosemi system with a sampling rate of 512 Hz. The subjects heard a single speech stream, in contrast to other studies (e.g. [O’Sullivan et al., 2015]) in which subjects were presented with two (or more) concurrent speech streams.

### 2.2 Classification Task

Stimulus-response models were evaluated using a classificaton task that involved deciding which of two candidate speech stream segments gave rise to a given EEG segment (match-vs-mismatch single-talker classification task). We chose this task, based on single-talker data, as it permits the analysis to focus on improving the stimulus-response models and decoding algorithms from a signal processing perspective rather than dealing with the cortical dynamics of attention that is encountered in the commonly used AAD task.

### 2.3 EEG and audio preprocessing

We employed the same preprocessing procedures as in [Wong et al., 2018]. In short, 50 Hz line noise and harmonics were filtered from the EEG using a boxcar (smoothing) filter kernel of duration 1/50 Hz. The data were then downsampled to 64 Hz using a resampling method based on the Fast Fourier Transform (FFT). To downsample, this method reduces the size of the FFT of the signal by truncating frequency components above the Nyquist frequency. An inverse FFT is then used to restore the signal to the time domain. The mean was removed from each EEG channel. EEG was then highpassed at 0.1 Hz using a 4th order forward-pass Butterworth filter for low frequency detrending. The joint diagonalization framework [de Cheveigné and Parra, 2014] was employed to remove eye artifacts in an automated fashion as described in [Wong et al., 2018], using FP3 and FP4 channels to detect eyeblink timepoints. For the the backward model, the EEG data was further bandpassed between 1-9 Hz using a windowed sync type I linear-phase finite-impulse-response (FIR) filter, shifted by its group delay to produce a zerophase [Widmann et al., 2015], with a conservatively chosen order of128 to minimize ringing effects. This frequency range was chosen as it has been shown that the cortical responses time-lock to speech envelopes in this range [O’Sullivan et al., 2015].

To obtain broadband audio envelopes, the presented speech stimuli were filtered into 31 frequency bands via a gammatone filterbank with a frequency range of 808000Hz [Patterson et al., 1987]. Each frequency band was fullwave rectified and raised to the power of 0.3 before being summed together. This step was intended to partially mimic the rectification and compression that is seen in the human auditory system [Plack et al., 2008]. The EEG and audio were subsequently downsampled to 64 Hz and aligned in time using start-trigger events recorded with the EEG. EEG channels and audio data were Z-normalized to their mean and standard deviation in the training data.

### 2.4 Cross-Validation Procedure

The classifiers described in the following sections were trained and evaluated on data for each subject using a 10-fold nested cross-validation procedure. This ensures that the test data used to evaluate the classifiers is not used during any part of the training process (including hyperparameter tuning). The data were divided into 10 folds and the outer cross-validation loop iterated over these folds. At each interation, 1 fold was held-out for testing, and the remaining 9 were used for training and hyperparameter tuning. Hyperparameters were tuned via an inner cross-validation loop: at each iteration of the inner loop, one fold was held out for validation and the remaining 8 were used for training. The objectives used for tuning the model hyperparameters are described with each model.

### 2.5 Stimulus-response models

Commonly-used stimulus-response models are shown in Figure 1. A forward stimulus-response model predicts the EEG from the speech envelope, a backward model infers the speech envelope from the EEG, and CCA maps both speech envelope and EEG data into a common subspace. Here we consider only backward and CCA-based models. The backward model, commonly used in decoding studies [Bialek et al., 1991, Mesgarani et al., 2009, Mesgarani and Chang, 2012, Ding and Simon, 2012b, Mirkovic et al., 2015, O’Sullivan et al., 2015, Van Eyndhoven et al., 2017, Wong et al., 2018], serves as a baseline by which other models can be evaluated. The title of the subsections describing each model (other than backward) or decoding scheme contains a code in brackets, to make it easier to refer to various combinations of these schemes.

**Figure 1:**
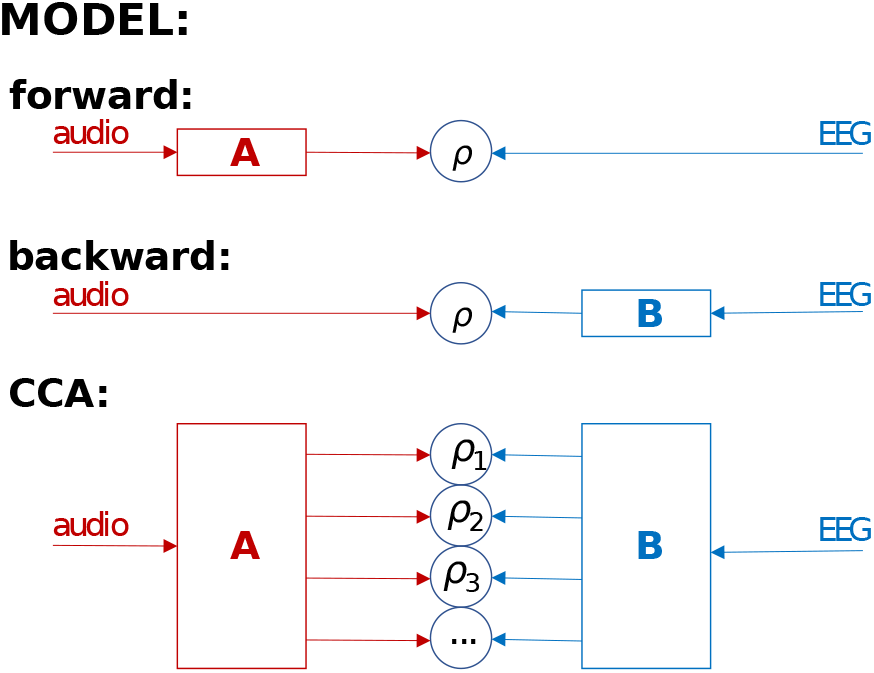
Three main stimulus-response models. The forward model predicts the EEG from the speech envelope. The backward model infers (“reconstructs”) the speech envelope from the EEG. CCA projects both speech envelope and EEG data onto components in a common subspace. Correlation coefficients (*ρ*) between predicted and actual EEG, inferred and actual stimulus, or canonical component (CC) pairs can be used as classification features.

#### 2.5.1 Data format and notation

The audio stimulus envelope is represented as a matrix **Y** = *y_t_* of size *T* × 1 where *T* is the number of samples. The EEG signal is represented as a matrix **X** = *x_t,n_* of size *T × N* where *N* is the number of channels. It may be useful to apply to each channel a set of *F* time shifts, or process the each channel by a *F*-channel filterbank. In that case **X** designates the resulting matrix of size *T × F N*.

#### 2.5.2 Backward Model

Backward models have been used extensively for the AAD [Akram et al., 2016, Mirkovic et al., 2015, 2016, O’Sullivan et al., 2015, 2017] and match-vs-mismatch classification tasks [de Cheveigné et al., 2018, Di Liberto et al., 2019]. The backward model has been shown to result in better classification accuracy than the forward model for these tasks, as it permits a spatial filter to be applied to the EEG to take advantage of inter-channel covariance to filter out brain signals unrelated to the auditory cortical response [Wong et al., 2018]. Here, we extend this scheme to permit a *spatiotemporal* filter by augmenting the EEG data by applying a set of time lags. Time lagged data are concatenated along the channel dimension to form a matrix **X** from which the audio envelope representation is inferred as **Ŷ**:

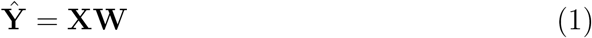

The weights **W** (spatiotemporal filter) are estimated from the training data using ridge regression as in [Crosse et al., 2015, 2016, Holdgraf et al., 2017, O’Sullivan et al., 2017, Wong et al., 2018]:

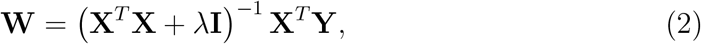

where λ is the regularization parameter and **I** is the identity matrix. The regularization parameter λ is optimized within the inner cross-validation loop to obtain the maximum correlation coefficient between the actual and predicted speech envelopes. An additional overall time shift parameter is also optimized within the inner loop. This time shift serves to absorb any latency mismatch due to filtering or cortical processing. The time-shift and λ parameters were optimized independently of each other, and in that order, for the purpose of saving time during model training.

#### 2.5.3 Canonical Correlation Analysis (C)

CCA finds linear transforms to apply to both audio and EEG to maximize mutual correlation. CCA has been shown to result in better classification accuracy than forward and backward models, as it allows spatiotemporal filters to be applied to both audio and EEG representations, stripping both of variance unrelated to the other [de Cheveigné et al., 2018]. CCA results in multiple pairs of canonical components (CCs), whereby the first has the largest correlation, and the second has the largest correlation that is orthogonal to the first, and so on. The audio and EEG CCs are computed as **C**_*Y*_ = **YW**_*Y*_ and **C**_*X*_ = **XW**_*X*_, respectively, where **Y** and **X** are the audio and EEG data, and **W**_*Y*_ and **W**_*X*_ are the corresponding spatio-temporal CCA weights.

Time lags can be applied to the EEG (as previously described for the backward model) as well as the audio representation (as typically applied in forward models) to allow the model to absorb convolutional mismatches between EEG and audio. However, to capture long-range temporal structure would require many lags, leading to computational issues and overfitting. For that reason, it is useful to replace the time lags by a smaller number of filters [de Cheveigné et al., 2018]. Here we use a set of *F*=9 dyadic filterbank kernels that approximate a logarithmic filterbank. The square-shaped left-aligned smoothing kernels have exponentially increasing lengths from 1 to 32 samples. The resulting audio data matrix **Y** has dimensions *T × F*, and the resulting EEG data **X** has dimensions *T × NF*, where the boxcar-smoothing and EEG channel dimensions are combined into a single dimension. Principal component analysis was applied to the filtered EEG data for whitening and regularization. For regularization, principal components beyond a certain rank were discarded before applying CCA. This is effectively a low rank approximation (LRA) regularization scheme [Marconato et al., 2014]. The optimal number ofEEG principal components to keep was determined as the number that maximized the cross-validated sum of correlation coefficients between CC pairs, over all pairs.

CCA was computed from the eigendecomposition of the covariance matrix **R** = ([**X, Y**]^*T*^[**X, Y**]), within the training dataset. The number of components, *n_cc_* is equal to the minimum size of the non-time dimension of **X** or **Y**. The CCA weights for **X, W**_*X*_, are contained within the NFx ncc upper-left sub-matrix in eig(**R**). Each column of **W**_*X*_ contains both channel and boxcar-smoothing dimensions, collapsed into a single dimension. The CCA weights for **Y, W**_*Y*_, are contained within the *F × n_cc_* lower-left sub-matrix in eig(**R**). An illustration of the CCA training inputs and outputs is shown in Figure 2.

**Figure 2:**
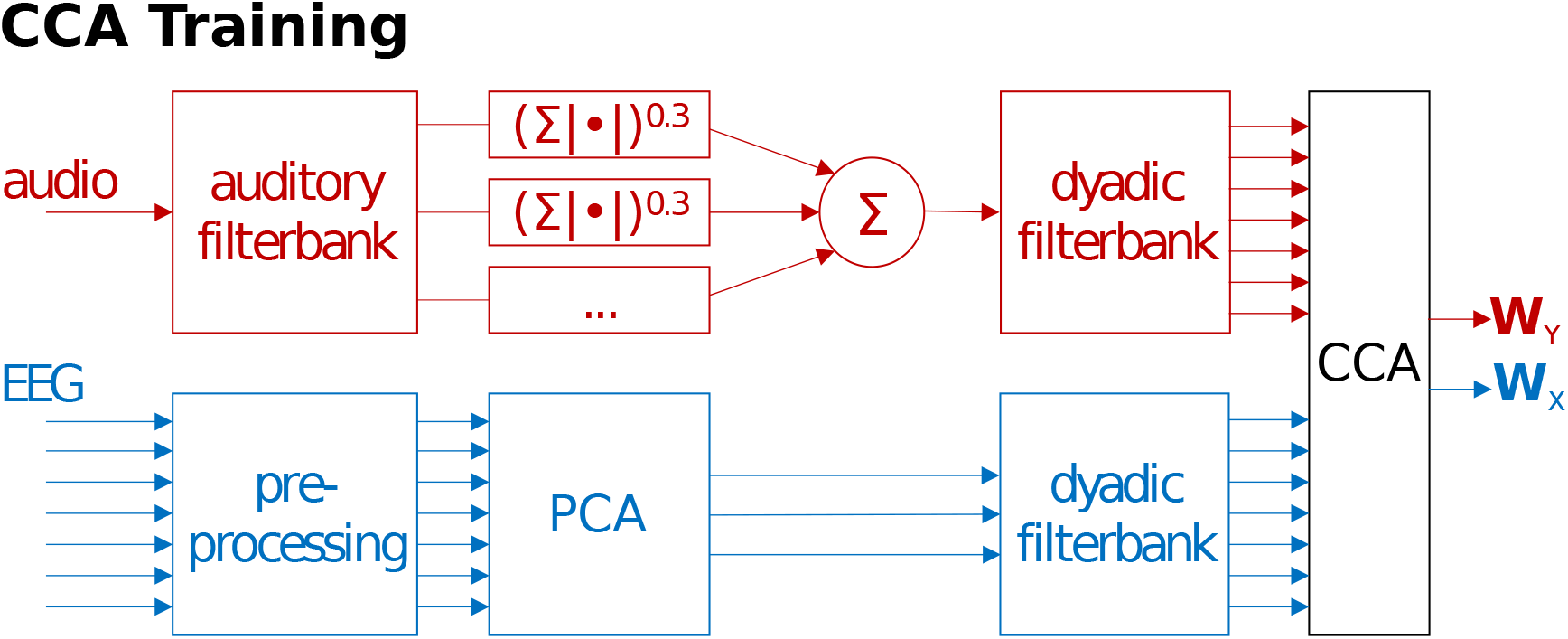
CCA training diagram. Preprocessed audio and EEG data are passed through a dyadic filterbank (see text). The CCA training algorithm then computes a set of weights **W**_*Y*_ and **W**_*X*_ that project the speech envelope and EEG data into a common subspace.

As for the backward model, an additional overall time shift parameter was introduced to absorb any temporal mismatch between stimulus and response due to filtering or cortical processing. This time-shift and the number of EEG principal components retained (see above) were determined within the inner cross-validation loop. They were determined independently and in that order to save computation. Classification schemes that involve the CCA model are indicated with a code that begins with the letter “C“.

### 2.6 Classification

The classification task is to decide, from a segment of EEG, which of two speech samples gave rise to it, the other being a sample from pseudorandom time window (“match vs mismatch” task). The features used for classification are, for the backward model, the correlation coefficient between the actual stimulus envelope and the estimate inferred from the EEG, and the correlation coefficient between the pseudorandom stimulus envelope and the estimate inferred from the EEG (bivariate feature). For the CCA model, the set of correlation coefficients between pairs of CCs is used (multivariate feature). The empirical joint distribution of features for matched and mismatched segments is estimated during the training phase of the classifier. For a new token containing an EEG segment paired with either the matching stimulus or a mismatching stimulus segment, the classifier identifies which of them corresponds to the match. Classification proceeds by situating the features relative to the empirical joint distribution for matched and mismatched pairs.

The classifier was trained anew on each iteration of the inner-cross-validation loop, using the model (backward or CCA) hyperparameters estimated on that iteration. The optimal hyperparameters and classifier found over iterations of the inner loop were then applied to classify data within the left-out fold of the current iteration of the outer cross-validation loop. The average of classification scores over iterations of the outer loop are reported in the Results. To generate classification data samples, the position of the decoding segment was stepped by 1s throughout the evaluated data. The *decoding segment duration* was chosen among values 3, 5, 7, 10 and 15s. These nominal durations include the length of the filtering kernels applied to the data (0.5s), as well as the optimal audio-EEG time-lag estimated in the hyperparameter estimation stage. Thus, they accurately reflect the duration of data used for each decision. The pseudorandom stimulus segment (foil) was drawn from a different fold from the actual speech sample. To allow reliable comparison between methods, the pseudorandom number generator was reinitialized with the same seed for the evaluation of each method.

For the backward model the classification feature was the correlation coefficient between the stimulus envelope and the envelope inferred from the EEG. To decode segment *d*, consisting of *D* time samples, the correlation coefficient between the predicted and actual speech envelope was computed as 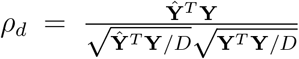. This feature was calculated for the stimulus segment within the test pair, and for a randomly chosen stimulus segment (foil). With this univariate feature, classification involves simply taking the larger correlation coefficient.

For the CCA model the classification feature was the set of correlation coefficients between selected CC pairs (9 pairs in the implementation presented here, since *F* = 9), as illustrated in Figure 3. This feature was calculated for the stimulus segment within the test pair, and for a randomly chosen stimulus segment (foil). These two multivariate values were fed to a multivariate classifier. We consider linear discriminant analysis (next section) to obtain baseline classification rates, and then proceed to neural network architectures.

**Figure 3:**
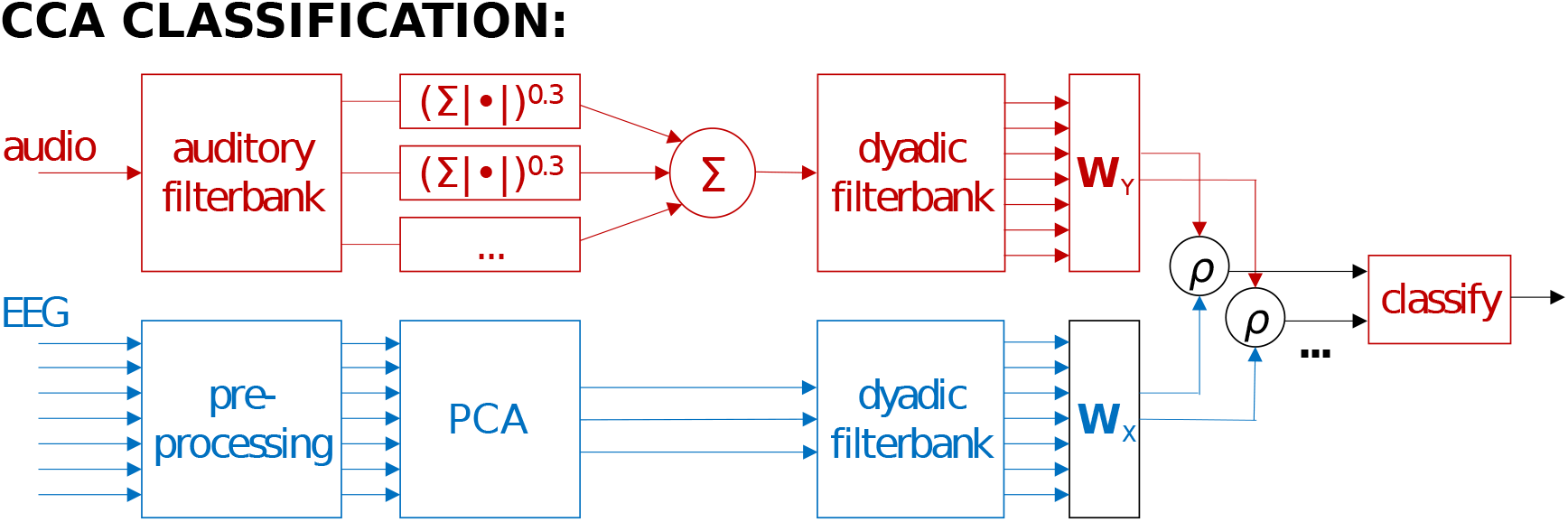
CCA classification diagram. Preprocessed audio and EEG data are passed through a dyadic temporal filterbank, then projected via weights **W**_*Y*_ and **W**_*X*_ learned by CCA onto CCs. Correlation coefficients computed over a decoding segment duration between CC pairs are used as features for classifying whether one of two audio streams is the one that corresponds to the EEG data, or comes from a random segment of speech.

#### 2.6.1 Linear Discriminant Analysis (L)

Denoting as *x_i_* the multivariate correlation coefficient feature (for consistency with standard expositions) and *y_i_* the class label for token *i*, the predicted class is computed as *ŷ_i_* = signum(*w ○ x_i_*), where *w* is a weight vector and the *y* ∈ {−1, +1}. LDA finds *w* such that the separation *S* between class distributions is maximized. *S* is defined as the ratio of the between class variance *σ_b_* to the within class variance *σ_w_*:

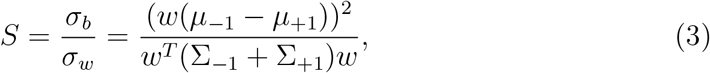

where *μ*_−1_ and *μ*_+1_ are the means of the two classes *x*_*i*|*y_i_*=−1_ and *x*_*i*|*y_i_*=+1_, and Σ_−1_ and Σ_+1_ are their standard deviations. *w* can be found by solving the generalized eigenvalue problem for the matrix 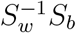, where *S_w_* is the within-class scatter matrix and *S_b_* is the between-class scatter matrix. Over all classes *c*, within-class scatter matrix is given by 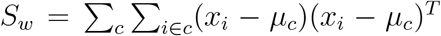. The between-class scatter matrix is given by 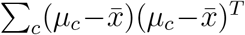. The eigenvector corresponding to the largest absolute eigenvalue is referred to as the first principal direction, or the weight vector *w*. The LDA classifier was trained on the correlation coefficients between the CCA-transformed audio (actual and random) and the EEG. A classification scheme that uses the LDA classifier is indicated by a code that ends in “*L*”.

### 2.7 Improving Classification Rates

The methods described so far correspond to those used in a previous paper that compared forward, backward and CCA models associated with LDA [de Cheveigné et al., 2018]. In this section we introduce several schemes that go beyond those methods with the aim of improving classification rates. These are of two sorts: the first two schemes aim at obtaining better linear transform weight matrices than those produced by CCA, the last three schemes make use of neural net architectures to make better use of the features for classification.

#### 2.7.1 D-Prime Maximization (D)

The cross-validation process described in Section 2.4 (inner loop) chooses hyperparameters so as to obtain the highest possible *sum of correlation coefficients* between CC pairs. Large correlation coefficients scores on matched pairs (compared to mismatch) might be expected to lead to good discrimination, however discrimination also depends on *intraclass variance* of those coefficients [Wong et al., 2018]. This is captured by the d-prime sensitivity metric, calculated as the ratio between the inter-class means and intra-class variance. At each iteration of the inner crossvalidation loop, a different set of CCA weights is computed for each regularization parameter sampled. By selecting those CCA weights that maximize the d-prime of output of a linear classifier applied to training data, classification error rates on the test set can potentially be reduced.

For each regularization parameter value sampled, the CCs computed on the validation data were divided into 2.5s segments. A classifier based on Kalman filtering was trained on the correlation coefficients between CC pairs for these segments. The derivation of this classifier permits a more analytic and stable evaluation of its d-prime score, although in practice it does not perform the match-vs-mismatch task as well as the LDA classifier. If we assume that the correlation coefficients between EEG and mismatching audio CCs have a mean of zero, and zero covariance with the correlation coefficients between EEG and matching audio CCs, the Kalman filter sensor matrix can be formulated as 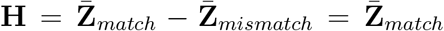 when the state *y* = 1 and 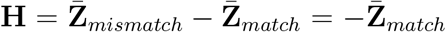 when the state *y* = −1 where **Z**_*match*_ = atanh(*ρ_match_*) and **Z**_*mismatch*_ = atanh(*ρ_mismatch_*). *ρ_match_* are the set of correlation coefficients between EEG and matching audio CCs. Similarly, *ρ_mismatch_* are the set of correlation coefficients between EEG and mismatching audio CCs. The hyperbolic-arctan-transform of is used to give **Z** a Gaussian distribution. The Kalman gain can then be written as 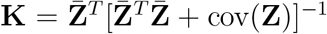, and the estimated states for each time sample in **Z** is then *ŷ* = tanh(**Z ∗ K**^*T*^), given a previous neutral state of 0. The d-prime for the classifier output is thus expressed as 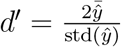. For simplicity, the same data used to train the Kalman classifier was used to compute the d-prime score.

Using an initial CCA regularization parameter value, an initial set of CCA weights was computed. The corresponding set of correlation coefficients between the resulting EEG and audio CCs, computed over 2.5s windows, was used to compute a Kalman classifier d-prime score. For each subsequent regularization parameter sampled, individual CCs were substituted into the previously established set and the d-prime score was recomputed. If an updated CC increased the d-prime score, the CCA weight corresponding to the updated CC was accepted as the new CCA weight. Abbreviated references to classification schemes implementing this method will include “*D*” in their name. For example, when CCA is applied using d-prime maximization and classification performed using LDA, this scheme will be denoted as “*CDL*”.

#### 2.7.2 Adaptive Beamforming (B)

Given a training data set, CCA produces a set of spatiotemporal weight matrices that optimize correlation between CC pairs on the training data. The EEG weight matrix has two characteristics: (a) it preserves the useful brain activity that underlies the correlation and (b) it suppresses sources of noise that would otherwise degrade that correlation. When the trained solution is applied to new data, however, the correlation structure of the noise may have changed so the solution is no longer optimal. The structure of the useful brain activity is less likely to change over time.

This situation can be addressed by applying a linearly constrained minimum variance (LCMV) beamformer. The LCMV beamformer, initially developed for antenna arrays, has proven useful to isolate localized neural activity by finding a weighted sum of EEG channels that project unit gain on a particular spatial location, while minimizing the contribution from all other locations. This type of beamforming is termed “adaptive” because the weights applied to the EEG channels are adjusted to minimize the noise based on the covariance structure *of the data being analyzed*. The LCMV beamformer requires knowledge of the forward model of the desired source (source-to-sensor matrix). This is usually assumed to require computation from knowledge of the source position, together with a geometric description of head tissues and tissue conductivity estimates, frequently taken from a structural MRI. However the formalism works just as well if the forward model is derived by other means. Here we derive it from the CCA solutions learned on the training set.

In this scenario, rather than corresponding to a specific spatial location, each “source” correspond to the forward model associated with a CC. Due to the orthogonal nature of the CCA weights, the mapping from the CCs to the EEG sensors is computed from **L** = [eig(**R**)^−1^]^T^, where **R** is the covariance matrix used to compute CCA from training data as described in Section 2.5.3. The first *n_cc_* columns and *NF* rows of **L** correspond to the forward potentials of the *n_cc_* CCs. We refer to this approach as “blind” in that it does not require knowledge of the actual geometry.

LCMV beamforming allows for the computation of weights that minimize noise within the EEG *test* data, and not just the training data. A forward model is derived from each of the CCs produced by applying CCA to the training data, based on which LCMV computes a beamforming weight vector that is used in lieu of the corresponding CC weight vector. In contrast to the CC weight vector that is fixed (after training) the beamforming weight vector is *adaptive*. This is useful in realtime applications where the nature of the noise is not always predictable, and also in batch processing of data with a complex non-uniform noise correlation structure.

In typical applications of LCMV to EEG data, such as neural source imaging, the forward potentials only contain a channel dimension, and sufficiently accurate forward potentials can be computed from a conductivity model of the head so that source localization can be performed. However, the CCA components yielded here contain both channel and boxcar-smoothing dimensions, combined into a single dimension. This larger dimensionality and the estimation of the source forward potentials from the data mean that these forward potentials are inexact. Errors in the forward potential can degrade beamformer performance, potentially resulting in the source of interest not being detected Dalal et al. [2014]. We use source suppression constraints to improve the solution, at the cost of reduced degrees of freedom for satisfying the beamforming objective of minimizing signal power. Given that each column in **L** is uncorrelated with each other, and to each CC being measured, this relationship can be enforced in the beamformer solution by introducing them as source suppression constraints Dalal et al. [2006], Wong and Gordon [2009].

The typical LCMV beamformer constraints are 1) enforce unit gain on the EEG source corresponding to a given CCA component and 2) minimize signal power. These constraints yield the following beamformer equation [Van Veen et al., 1997]:

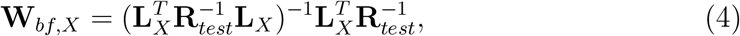

where **R**_*test*_ is the data covariance matrix computed in a similar way to **R**, but over the validation or test fold. **L**_*X*_ is the CCA forward potential column-vector, computed from the training data. Given that inaccuracies in the CCA forward potential estimate would result in reduced SNR and leakage from other sources, we add an additional constraint 3) enforce nulls on the EEG sources corresponding to *S* uncorrelated sources, where *S* is optimized by cross-validation. This effectively minimizes the contribution of noise leakage into the beamformed signal. These uncorrelated source constraints are drawn from other columns in **L**, which are orthogonal by definition. This third constraint is implemented by structuring **L**_*X*_ such that the first column is the forward potential corresponding to an individual CC being measured, and the remaining columns are the forward potentials of the sources to be suppressed. These columns are taken from the *N F × S* upper-left sub-matrix in **L**.

Given that a larger number of suppression constraints reduces the degrees of freedom available to the beamformer to suppress noise sources, the optimal number of suppression constraints *S* needs to be determined. This is done via the 9-fold inner cross-validation described in Section 2.4. *S* was determined separately for each CC. Thus, the beamforming implementation with CCA effectively involves tuning three types of regularization parameters to maximize the cross-validated sum of correlation coefficients across CC pairs: the number of lags, the number of principal components kept when whitening EEG, and the number of suppression constraints per CC. The number of lags is determined first, independent of the others. The number of principal components to keep and the number of suppression constraints are then determined via a grid search. Note that here as with the default CCA implementation, the same number of principal components is kept for all CCs. Classification schemes implementing this method will include “*B*“ in their name.

Beamforming and d-prime maximization can be combined. With d-prime maximization and no beamforming, while adjusting the number of principal components kept during EEG whitening as a regularization parameter, individual CCA weights that maximized the validation classifier d-prime were kept. When combined with beamforming, rather than keeping the individual CCA weights, the individual CCA forward potentials and associated source suppression constraints are kept instead.

#### 2.7.3 Multilayer Perceptron (M)

The LDA classifier uses only the principal direction in multivariate space to separate the two classes. Other directions, possibly also informative for class separation, are ignored. A multilayer perceptron (MLP) neural network can find a nonlinear decision function that may be better as it combines information from multiple decision planes. We implemented a multilayer perceptron (MLP) neural network with hyperbolic tangent activation functions, feeding into a two-unit softmax classification layer. An MLP layer performs a nonlinear operation on the inputs *y_i_* = tanh(**W***x_i_ + b*), where **W** is the weight matrix and *b* is the bias vector. Multiple MLP layers can be stacked so that subsequent layers take the output from the previous layer as input. A softmax layer takes the output from the last MLP layer as its input and computes an output such that each value *c* in *y_i_*, corresponding to each class, is normalized according to 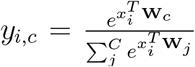. The largest of the values *c* in *y_i_* corresponds to the predicted class. The network was trained using a categorical cross-entropy cost function. Training was performed using minibatches, and *rmsprop* as the gradient descent method [Hinton et al., 2012]. Early stopping was used to terminate training when the validation cost function no longer improved. Different numbers of MLP layers and units per layer were experimented with. Abbreviated references to the CCA classification schemes using an MLP will end in “*M*”.

#### 2.7.4 Simple Recurrent Layer (S)

Up to this point, single correlation coefficients have been computed over the entire segment of data used for classification. A correlation coefficient is the dot product between two normalized CC time series in which all time points are weighted equally. However, if information could be obtained as to which time points are more reliable, it would be more appropriate to apply a non-uniform weight to modulate the amount each time point contributes to the final classification. We divided the correlation coefficient computation over the entire decoding segment into non-overlapping subintervals of one second duration. The number of sub-intervals was thus equal to the decoding segment duration in seconds. Based on the correlation coefficient equation, the *sub-interval correlation coefficient* computed over a sub-interval *s* is computed as:

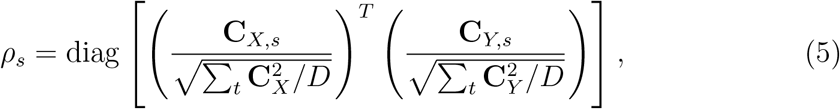

where for a total of *S* sub-intervals, **C**_*X,s*_ ≡ **C**_*X*_(*t* ∈ (*s, s* + 1]*D/S*), and similarly **C**_*Y,s*_ ≡ **C**_*Y*_(*t* ∈ (*s, s* + 1]*D/S*). This denominators of this equation are computed over the entire decoding segment duration, whereas the numerators are computed only over the sub-interval. Since the denominator can be seen as a normalization factor, computing *ρ_s_* in this way stabilizes the normalization factor over the entire interval.

To determine the weighting for each sub-interval, we chose to employ a simple recurrent network (SRN) layer which takes only the sub-interval correlation coefficients as inputs. An SRN takes a set of input vectors over *S*-time steps, *x_s_*. For each time step, it computes a new state *h_s_* based on its previous state *h*_*s*−1_ and input vector *x_s_* according to *h_s_* = tanh(**W**_*h*_*x_s_*+**U**_*h*_*h*_*s*−1_+*b_h_*). The output ofthe last SRN timestep was passed to a 2-layer MLP, consisting of 3 units each, terminating in a softmax output layer for classification. Training was performed using minibatches, and *rmsprop* as the gradient descent method [Hinton et al., 2012]. Abbreviated references to the CCA classification scheme using an SRN will end with “*S*”.

#### 2.7.5 Gated Recurrent Unit (G)

An SRN lacks the ability to store information over long durations due to the vanishing gradient problem: the SRN time-steps are unfolded into a multi-layer network for training, and with the use of sigmoid-like activation functions, the backpropagated error diminishes across layers, preventing the SRN from learning long-term relationships [Pascanu et al., 2013]. In contrast, a gated recurrent unit (GRU) allows the error to be preserved through time and layers. A GRU updates its internal state *h_s_* based on two gating functions: the update gate *z_s_* and the reset gate *r_t_* [Cho et al., 2014]. The update gate determines how much of the current state *h_s_* at timestep *s* incorporates the previous state *h*_*s*−1_ versus a candidate state *h̃_s_* computed from *x_s_* plus some leakage from *h*_*s*−1_.

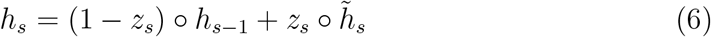

The update gate *z_t_* is computed as a sigmoid function of the weighted GRU input *x_t_* and the previous state *h*_*t*−1_:

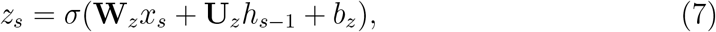

where **W**_*z*_ and **U**_*z*_ are weights and *b_z_* is a bias vector. The candidate state *h_s_* is computed as a hyperbolic tangent function of the weighted GRU input *x_s_* and the weighed previous state *h*_*s*−1_:

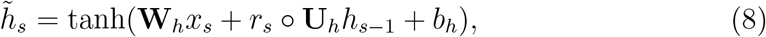

where **W**_*h*_ and **U**_*h*_ are weights and *b_h_* is a bias vector. The reset gate, *r_s_* determines how much leakage from the previous state is incorporated into *h_s_*. Similar to the update gate, is computed as a function of weighted GRU input and the previous state.

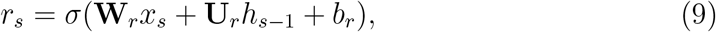

where **W**_*r*_ and **U**_*r*_ are weights and *b_r_* is a bias vector.

The GRU layer consisted of 8 units. The output of the last GRU timestep was passed to a 2-layer MLP, consisting of 3 units each, terminating in a softmax output layer for classification. Abbreviated references to the CCA classification scheme using an GRU will end with “*G*“.

### 2.8 Classifier Performance Evaluation

We used two metrics to quantify performance: classification error rate and information transfer rate (ITR). The ITR is the number of correct decisions that can be made by the classifier per unit time. Because increased decoding segment lengths result in a reduction in the number of decisions that can be made per unit time, this measure allows for the comparison of results across different decoding segment lengths. The ITR measure that was used was the Wolpaw ITR [Wolpaw and Ramoser, 1998] and is calculated by:

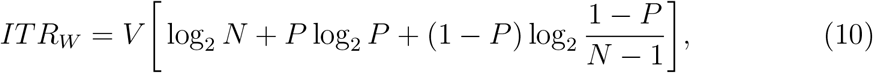

where *V* is the speed in trials per minute, *N* is the number of classes, and *P* is the classifier accuracy (1 minus the error rate). Both metrics were averaged across all test folds for each subject.

### 2.9 Implementation

Data preprocessing and CCA analyses were performed using the COCOHA Matlab Toolbox [Wong et al., 2018]. The scikit-learn implementation of LDA was used, and the neural networks were implemented in Keras [Chollet et al., 2015].

## 3 Results

A summary of the classification performance of all methods is shown in Figure 4. Performance is quantified here as percent error rate rather than percent correct rate as is common: lower is better. The left panel shows the average error rate over subjects for a range of decoding segment lengths, and the right panel shows the error rate at a 5s decoding segment length for each subject. Moving left to right, a clear improvement can be seen as new methods are introduced and combined. Taking the backward model as a baseline, the best combination reduces the error rate by from 18.9% to 3.0% (i.e. by a factor of 6.3). The contribution of each step is detailed in the following. For paired t-test analyses of error rate data, a logit transform is applied to the error rates [Warton and Hui, 2011].

**Figure 4:**
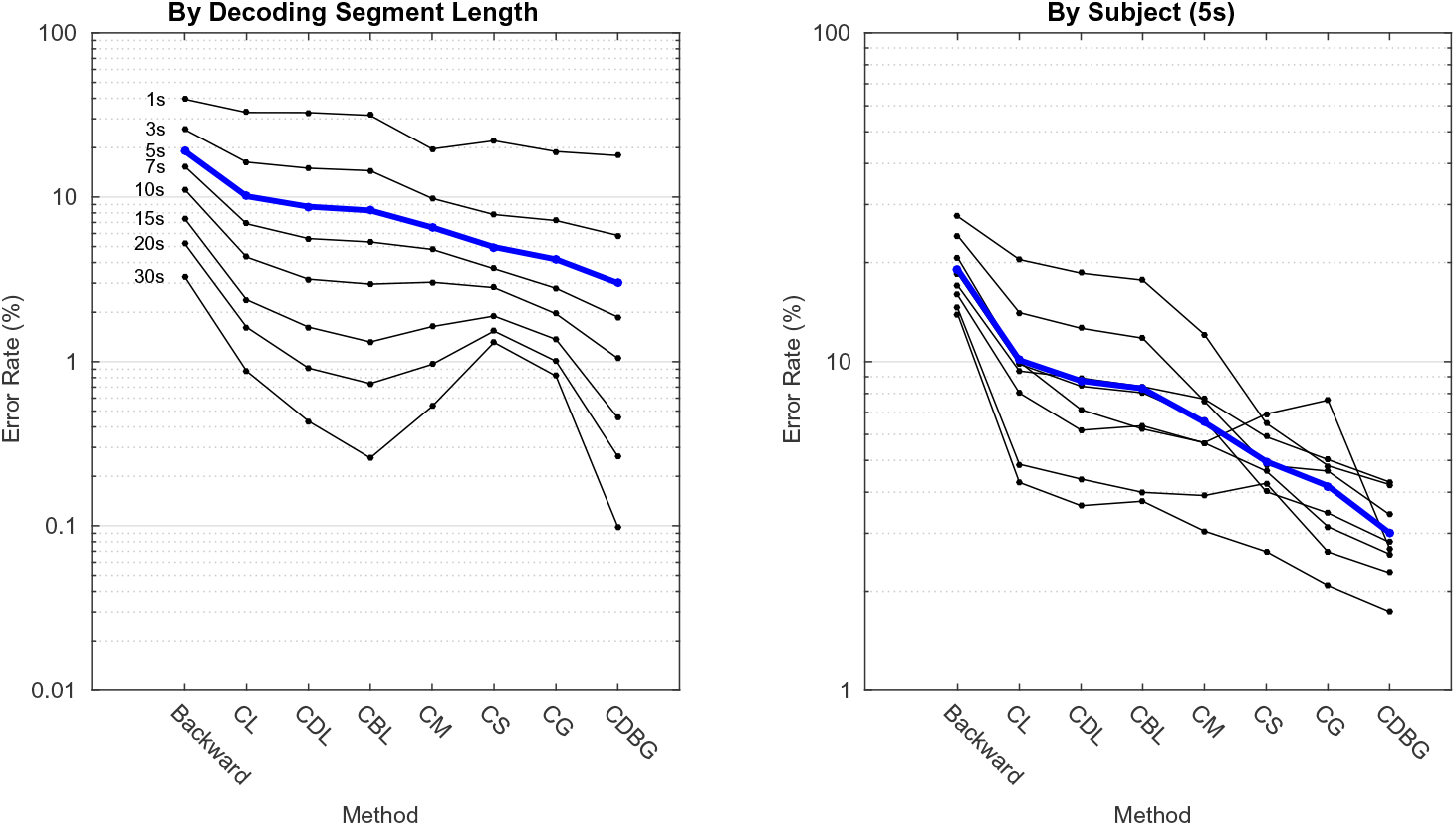
Classification error rate for different classification schemes. Chance is 50%. The left panel shows error rates with different decoding segment lengths, averaged over all subjects. The right panel shows error rates for each subject using a 5s decoding segment length. The average of all subjects is indicated by the blue line, which also corresponds to the blue line in the left panel. *C* = CCA, *D* = d-prime maximization, *B* = beamforming, *L* = LDA, *M* = MLP, *S* = SRN, *G* = GRU.

### 3.1 CCA vs backward model

CCA+LDA (*CL*) provides a clear improvement over the backward model, as we found previously [de Cheveigné et al., 2018]. At a decoding segment length of 5s the error rate decreased by 9.0 percentage points (paired samples t-test, *T*_79_ = 21.9, *p* = 2.7 × 10^−35^), that is by a factor of1.89, on average over subjects. The difference is of same sign for all subjects, and all durations. It is instructive to see how this improvement relates to the original error rate.

Figure 5 left shows a scatterplot of error rates for the CCA+LDA scheme vs the backward model. Each dot represents the error rate for one test fold and one subject. The axes are scaled by a logit transform to account for the saturation effect as the error rate decreases to 0 [Warton and Hui, 2011]. This transform produces a normal distribution and equivariance in regression residuals, which are underlying assumptions of linear regression model statistics. On these axes the data follow a linear trend with slope *m* = 1.47 greater than one (*CI*_.95_ = [1.29, 1.66]). This indicates that the benefit was greater for classification folds that already had a low error rate, after accounting for the effects of saturation.

We now use the CCA+LDA model (*CL*) as a baseline to evaluate schemes that further improve performance. We report the effect of scheme is shown in isolation (relative to *CL*) as well as their best-performing combination (*CDBG*). We also analyze improvement as a function of the baseline error rate, as summarized in Figure 5 (center).

**Figure 5.**
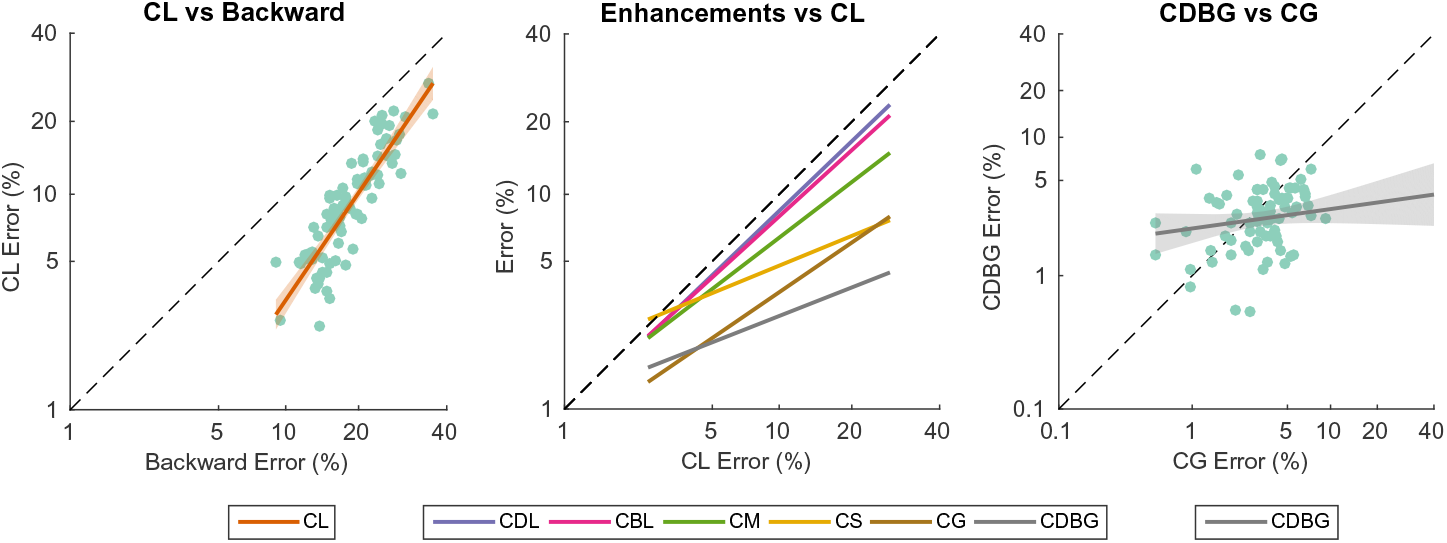
Error rate relationships between classification schemes, with lines of best fit. The graphs are plotted on logit axes in order to compensate for effects of saturation as error rates approach 0 or 100% [Warton and Hui, 2011]. The left panel shows the error rate of CCA+LDA (*CL*) versus that of the backward model. The center panel shows linear fits to scatter plots of error rates of several enhanced schemes relative to CCA+LDA (*CL*). The right panel shows a scatterplot of the error rate for the best combination (CDBG) relative to the second best (*CG*). Dots in left and right panels represent classification errors for test folds, shown for all subjects. Translucent bands around the lines in the left and right panels indicate the 95% confidence intervals. *C* = CCA, *D* = d-prime maximization, *B* = beamforming, *M* = MLP, S = SRN, *G* = GRU.

### 3.2 D-Prime Maximization (D)

D-prime maximization (see Methods) yielded a 1.4 percentage point (a factor of 1.16) classification error decrease (paired samples t-test, *T*_79_ = 5.9, *p* = 6.6 × 10^−8^). The purple line in Figure 5 (center) represents a linear fit of the scatter plot error rates of *CDL* vs *CL*. The slope *m* = 0.96 was not significantly different from 1. In other words, maximizing the d-prime scores equally reduces the classification error of all folds regardless of the original CCA+LDA classifier (*CL*) error.

### 3.3 Beamforming (B)

Beamforming (see Methods) yielded a 1.8 percentage point (a factor of 1.22) classification error decrease (paired samples t-test, *T*_79_ = 7.1, *p* = 4.8 × 10^−10^). The red line in Figure 5 (center) represents a linear fit to the scatterplot of error rates of (*CBL*) versus *CL*. The slope *m* = 0.91 was not significantly different from 1.

### 3.4 Multilayer Perceptron (M)

A four-layer MLP network, with 8 units in the first layer and 3 units second and third layers, followed by a 2-unit softmax classification layer, achieved an 3.6 percentage point (a factor of 1.55) classification error decrease over the original CCA+LDA classifier (*CL*) (paired samples t-test, *T*_79_ = 12.3, *p* = 5.2 × 10^−20^). The green line in 5 (center) represents a linear fit of the scatterplot of error rates of *CM* vs *CL*. The slope of *m* = 0.77 was significantly less than 1 (*CI*_.95_ = [0.66, 0.88]), indicating that the error rate decreased more for folds that had larger error rates. Increasing the number of layers or units per layer did not significantly impact the classification performance. Simple Recurrent Network (S)

### 3.5 Simple Recurrent Network (*S*)

Replacing the first MLP layer with an 8-unit simple recurrent network (SRN) achieved a 5.2 percentage point (a factor of2.04) classification error decrease (paired samples t-test, *T*_79_ = 11.9, *p* = 2.4 × 10^−19^). The yellow line in Figure 5 (center) represents a linear fit of the scatterplot of error rates of *CS* vs *CL*. The slope *m* = 0.41 was significantly less than 1 (*CI*_.95_ = [0.26, 0.56]), indicating that the error decreased more for folds that had larger error rates.

It is worth noting that MLP and SRN classifiers perform less well at longer durations (Figure 4, left), and at 20 and 30s the SRN classifier (*CS*) yields greater error rates the original *CL* scheme. This is possibly a result of the vanishing gradient problem which prevents the SRN from learning long-term relationships, and thereby impedes performance when the recurrent classifier must make a prediction after processing a larger number of sub-intervals.

### 3.6 Gated Recurrent Unit (*G*)

Replacing the SRN layer by an 8 unit GRU layer yielded a 5.9 percentage point (a factor of 2.42) classification error decrease (paired samples t-test, *T*_79_ = 15.8, *p* = 4.3 × 10^−26^). The brown line in Fig. 5 (center) represents a linear fit of the scatterplot of error rates of *CG* vs *CL*. The slope *m* = 0.68 was significantly less than 1 (*CI*_.95_ = [0.47, 0.89]). Again this indicates that the error decreased more for folds that had larger error rates, although folds with small error rates also seem to benefit (Figure 5 center).

The classifier with a GRU layer (*CG*) performed better than a classifer with a SRN layer (*CG*, (paired samples t-test, *T*_79_ = 4.4, *p* = 3.8 × 10^−5^). To determine whether this could be due to the larger number of parameters used in the GRU network (693 parameters, including the MLP portion), we implemented also a classifier with an SRN layer with 17 units (684 parameters). The classification error for the larger SRN was significantly larger than that obtained by the 8-unit SRN by 1.3 percentage points (paired samples t-test, *T*_79_ = 7.2, *p* = 3.1 × 10^−10^) suggesting overfitting. The advantage of the GRU is thus unlikely to be related to its larger number of parameters.

### 3.7 Combined Methods (*CDBG*)

The GRU (*CG*) yielded the largest decrease in error rate over the basic CCA+LRA implementation for durations up to 10s (Figure 4 left). However, combining it with several of those schemes yielded a yet greater improvement (*CDBG*). Adding d-prime maximization and beamforming reduced the error rate by 1.2 percentage points (paired samples t-test, *T*_79_ = 2.49, *p* = 0.015), that is a factor of 1.39. Interestingly, this benefit extended also to long durations (Figure 4 left), attaining an error rate of 0.1% for 30s duration segments (compared to 3% for the backward model).

Figure 5 (right) shows a scatterplot of error rates for CDBG relative to *CG*. The slope *m* = 0.14 is significantly smaller than 1 (*CI*_.95_ = [−0.04, 0.34]), indicating that the improvement is greatest for folds/subjects for which error rates were relatively high.

### 3.8 Information Transfer Rate (ITR)

From Figure 4 (left) it is obvious that there is a tradeoff between error rate and segment duration, shorter segments yielding greater error rates. An alternative metric of performance is ITR (roughly, the number of decisions that can be made per unit time, see Methods). Such a metric is relevant for BCI applications that require decisions to be both accurate and timely.

Figure 6 plots values of the ITR for the backward model (red), CCA+LDA (*CL*, blue), and CCA with d-prime maximization, beamforming, and GRU neural network improvements (*CDBG*, green). As expected from the error rate metric, the more sophisticated schemes yield higher ITR rates. The maximum ITR is reached at 5s for the backward model, 3s for *CL* and 1s for *CDBG*).

**Figure 6:**
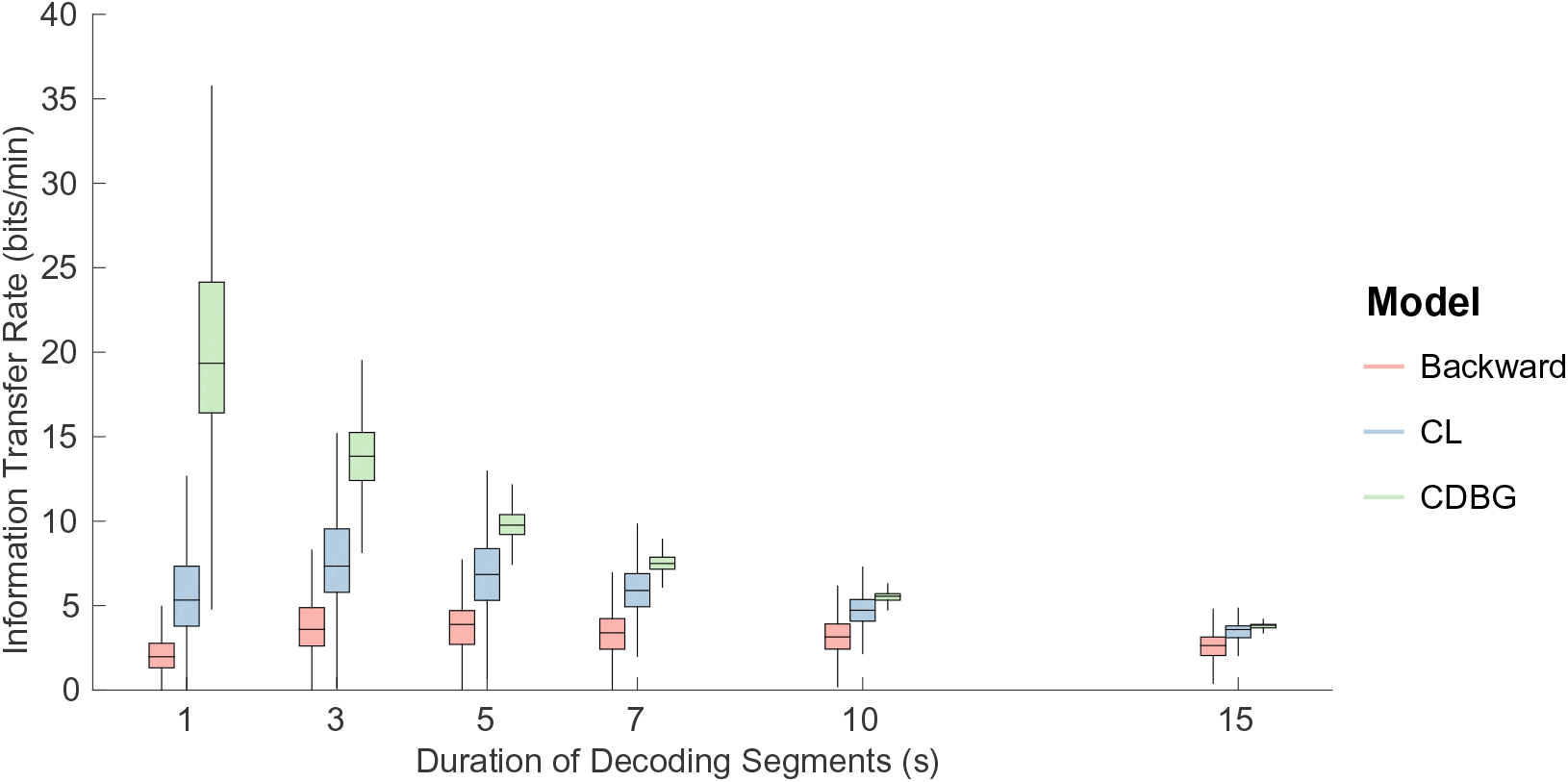
Information transfer rate comparison over different decoding segment durations for the backward model, the baseline CCA implementation using an LRA classifier (*CL*), and the CCA implementation with all enhancements: d-prime maximization, beamforming and a GRU classifier (*CDBG*).

## 4 Discussion

In previous work we found that a CCA-based model yielded more accurate classification than standard backward or forward models [de Cheveigné et al., 2018], presumably thanks to the ability of CCA to factor out irrelevant information from both audio and EEG, and to provide multiple components to support multivariate classification. In the present paper, we observed that the benefit over the backward model (in relative terms) was smaller for folds where the backward model gave larger classification errors (Figure 5 left), suggesting that performance might be limited by poor EEG data quality on those folds. Thus, we focused on improving the CCA classification framework to be more robust to noise (defined as any feature of the data that increases classification error). This encompasses EEG artifacts, but it might also include points in the audio that do not yield reliable EEG response, such as silences. The solutions explored were methods to improve estimation of the CCA weights, and to allow a classifier to utilize temporal information.

### 4.1 Improving CCA Weights

When applied to CCA+LDA, both d-prime maximization and beamforming reduced classification errors equally across classification folds, regardless of the original error. Maximization of component d-prime yielded EEG spatial filter weights that were superior to those provided by CCA. This operation was performed individually for each CC. An alternative approach, not explored in the present study, could be to maximize the d-prime output or the loss function of a classifier via tuning of all components in combination.

Beamforming is another approach to improve spatial filter weights. It requires knowledge of the forward potentials of sources to preserve. Typically this knowledge is computed from anatomical data and models of head tissue conductivity, but here we use forward potentials associated with optimal components computed from CCA. Beamforming adaptively suppresses activity other than that associated with the forward potentials, effectively addressing the time-varying structure of the noise. We did not make full use of this flexibility in our simulations: beamforming was applied on the basis of the covariance matrix calculated over the full length of the cross-validation fold, which is roughly 9 minutes. An alternative, not explored in the present study, is to recalculate the beamformer solution based on shorter intervals. There is, however, a limit to which the time window can be shortened as sufficient data is needed to accurately estimate the covariance matrix **R**.

### 4.2 Improving the Classifier

A multilayer perceptron (MLP) network reduced classification errors slightly compared to an LDA classifier, suggesting that there is some advantage that can be gained from a nonlinear decision function. However, the recurrent neural networks (SRN and GRU) showed the largest reduction in classification error over an LDA classifier. The recurrent networks yielded the greatest benefit for folds with higher CCA+LDA classification errors, suggesting that they can tackle noise features for which the other classifiers fail. The recurrent layers are likely able to handle shorter-term variations in the noise, compared to d-prime maximization or beamforming, that are calculated over the entire cross-validation/test dataset. The time-scale of variations in the noise that can be handled by the SRN or GRU are related to the length of the sub-intervals used to compute the correlation coefficients fed to these neural network layers. While the GRU provided the largest reduction in classification error over CCA+LDA, combining it with component d-prime maximization and beamforming provided a significant additional reduction.

### 4.3 Relation between same-different and AAD tasks

The results reported in this paper were obtained for a match-vs-mismatch classification task, that allowed us to focus on the quality of the stimulus-response model. We preferred this task to the more complex AAD task, as it is not vulnerable to mislabeling of the database. In the AAD task an “error” might be the result of attention drift, making it hard to explore the performance in the region of low error rates (of use for applications). Cortical responses to concurrent speakers have been shown to have slightly different dynamics than those to a single speaker. [Ding and Simon, 2012b] found that the attended talker shows a stronger representation than the unattended talker at longer latencies (≈200ms), whereas both attended and unattended talkers are equally represented at shorter latencies (≈80ms). We expect our methods to be effective also in the AAD task, but it would be useful to confirm this in future studies.

Extrapolating from our results, and considering the many paths that remain to be explored, we believe that further improvements in classification performance may be possible.

### 4.4 Summary

Previous studies showed that the relation between stimulus and brain response can be captured by a linear model fit using system identification techniques, extending classic ERP studies to allow continuous stimuli such as speech [Lalor et al., 2006, 2009, Lalor and Foxe, 2010, Power et al., 2012]. Such a linear model can be used by a classifer in a BCI application, for example to decide whether a listener is attending to one or the other of two concurrent voices (AAD), but poor classification reliability and the amount of data required by each decision limit the practical use of such a scheme [O’Sullivan et al., 2017, Zink et al., 2017, Wong et al., 2018]. In previous work we showed that the stimulus-response model can be significantly improved using CCA [de Cheveigné et al., 2018], and here we showed that classification performance can be further enhanced by improving the quality of EEG linear filters over CCA, or improving the classifier over LDA. Overall, the error rate was divided by 6 over the standard backward model, for a 5s segment of data. This brings us closer to the goal of reliable “cognitive control” of a device based on brain responses.

## 5 Acknowledgements

This work was supported by the EU H2020-ICT grant 644732 (COCOHA), and grants ANR-10-LABX-0087 IEC and ANR-10-IDEX-0001-02 PSL. It draws on work performed at the 2016 Telluride Neuromorphic Engineering workshop.

